# Assays for monitoring *Toxoplasma gondii* infectivity in the laboratory mouse

**DOI:** 10.1101/584474

**Authors:** Qiuling Wang, L. David Sibley

## Abstract

*Toxoplasma* is a widespread parasite of animals including many rodents that are a natural part of the transmission cycle between cats, which serve as the definitive host. Although wild rodents, including house mice, are relatively resistant, laboratory mice are highly susceptible to infection. As such, laboratory mice and have been used to compare pathogenesis of natural variants, and to evaluate the contributions of both host and parasite genes to infection. Protocols are provided here for evaluating acute and chronic infection with different parasite strains in laboratory mice. These protocols should provide uniform standards for evaluating natural variants and attenuated mutants and for comparing outcomes across different studies and between different laboratories.

## 1 Introduction

*Toxoplasma gondii* is an extremely widespread parasite of animals that also causes zoonotic infections in humans (1). Strains of *T. gondii* have been grouped into three major clonal lineages that predominate in North America and Europe (2–4). These lineages differ in their acute virulence in laboratory mice due to the presence of polymorphic secretory effectors, many of which are derived from rhoptry secretion (5) or released from dense granules (6). Representatives of all three genotypes have been reported as part of a comparative genomes project for *T. gondii*, which provides a framework for comparing strain types within and between major lineages (7). Strains of *T. gondii* from South America, which differ genetically, also show high levels of virulence in laboratory mice, in part due to similar virulence factors that mediate differences among the clonal lineages (8). Here we will present protocols for monitoring infectivity and pathogenesis of the three major clonal lineages in laboratory mice, although similar methods could be adapted to study other more diverse strains.

Transmission of *T. gondii* normally occurs between cats, which serve as the definitive host, and rodents and many other animals that serve as intermediate hosts (1). Mice are a natural host for *T. gondii* and as such the laboratory mouse provides an excellent model to study innate and adaptive immunity (9, 10). Type I strains are acutely virulent and a single viable organism is uniformly lethal in all strains of laboratory mice (11, 12). Type I strains do not readily differentiate to bradyzoites in mice and their high level of virulence makes it difficult to obtain tissue cysts in chronically infected animals. As a consequence, infections with Type I strains are typically administered by intraperitoneal (IP) injection of tachyzoites. Acute infections progress rapidly by expansion of parasite numbers and dissemination to all organs of the body, leading to death within the first 10-12 days (13). Parasite expansion and dissemination are prominent features of acute infection, leading to cytokine shock (13, 14), which is likely a contributing cause of death. Genetic crosses have been used to map genes that contribute to acute virulence of Type I strains in laboratory mice (15) and the roles of specific genes in pathogenesis have been confirmed using a variety of techniques to disrupt or modify genes (16).

By contrast, Type II strains display intermediate virulence in laboratory mice where high doses lead to lethal outcome, while lower doses resolve and result in chronic infection, allowing LD_50_ values to be established (13). Because of their ability to cause nonlethal, chronic infection, Type II strains are often used to explore a range of immunological functions that control infection (9, 10). Infection protocols vary with some investigators using IP injection of tachyzoites grown in vitro, while others isolate tissue cysts from chronically infected mice and administer them by IP injection or by oral gavage. Oral ingestions of tissue cysts follow the natural route of infection as *T. gondii* is able to transmit between different intermediate hosts by omnivorous or carnivorous feeding (17). High challenge doses delivered by the oral route result in acute gastroenteritis that can lead to death, and the immunological basis of this form of pathogenesis has been explored through numerous studies (18).

Type III strains are relatively common in animals in North America and yet they are rarely encountered in human cases of toxoplasmosis (4, 19, 20). Type III strains are highly avirulent in laboratory mice, with high challenge doses leading to low levels of lethal infection (21). The Type III strain CTG (aka CEP) has been used to study sexual phase transmission in the cat and to develop genetic mapping strategies for *T. gondii* (22, 23). The basis for the lack of acute virulence in CTG was shown to be due to under-expression of ROP18, a polymorphic virulence determinant of Type I strains (24). Infection studies with another commonly used Type III strain called VEG revealed that the stage used for infection (i.e. bradyzoite vs. oocyst challenge) greatly influences pathogenesis (25).

Laboratory mice are derived from a few founder lines that represent mixtures of the *Mus musculis musculis*, *M. m. domestica*, and *M. m. castaneus* lineages, with the majority of loci coming from *M. m. domesticus* (26). These founders were used to establish outbred Swiss Webster and CD-1 lines, which have been kept in closed colonies and bred to maximize genetic heterozygosity (27). Inbreeding of founder lineages to minimize heterozygosity gave rise to C57Bl/6 mice and other inbred lines that differ by a small number of polymorphic loci, notably the major histocompatibility complex (MHC). Such inbred lines have been extremely useful for studying the association of genotype with phenotype. However the total variation within all inbred laboratory mouse lines is far less than that seen in wild caught or wild-derived isolates of *M. musculis* (28). Although both outbred and inbred strains of laboratory mice are relatively susceptible to *T. gondii* infection, wild strains derived from *M. musculis* are much more resistant (29). Given their susceptibility to infection, laboratory mice have been extremely useful to highlight pathogenesis differences among parasite strains and to study immune mechanisms involved in control of infection.

## 2 Materials

### 2.1 Propagation of tachyzoite cultures in vitro

1. Commonly used strains for mouse challenge studies are shown in Table 1. For type I strains we recommend the lab-adapted RH strain, or GT-1 which undergoes the complete life cycle. Commonly used type II strains include ME49 and Prugniaud (aka Pru). Type III strains include CTG and VEG. Estimates of the pathogenicity in different mouse strains are also provided, although these can vary with colony and source and so need to be determined locally.
2. Tissue culture incubator for culture at 37 °C, 5% CO_2_ and BSL-2 biosafety cabinet.
3. Tissue-culture flasks (25 cm^2^), 96-well plates, and cell scrapers.
4. Human foreskin fibroblasts (HFF) cells (ATCC, cat. # SCRC-1041).
5. D10 medium: For 1 liter, combine 1 package Dulbecco’s Modified Eagle’s Medium (DMEM) powder (Thermo Fisher, Gibco cat. # 12100046), 3.7 g NaHCO_3_, 100 ml fetal bovine serum (FBS), 10 ml of 200 mM L-glutamine, 1 ml of 10 mg/ml gentamycin, and dH_2_O to 1 liter. Sterilize by 0.45 micron filtration.
6. HHE: 1X Hank’s Balanced Salt Solution, 10 mM HEPES, 1 mM EGTA, sterilized by 0.45 micron filtration.
7. Blunt-end needles (20, 23, and 25 gauge), bulb transfer pipets, 5 and 10 ml pipets, 5 and 10 ml syringes.
8. Hemocytometer (Thermo Fisher Scientific, cat. # 0267151B) or other cell counting instrument.
9. Inverted tissue culture microscope equipped with 10x, 20x objectives.
10. Swin-Lok filter holder (Whatman, cat. # 420200) and polycarbonate filter membranes (3 micron pore) (Whatman, cat. # 110612). Although larger pore sizes can be used, there is more risk of contaminating host nuclei or debris.
11. Centrifuge capable of holding 15 conical tubes and spinning at 400 × *g.*

**Table 1.**
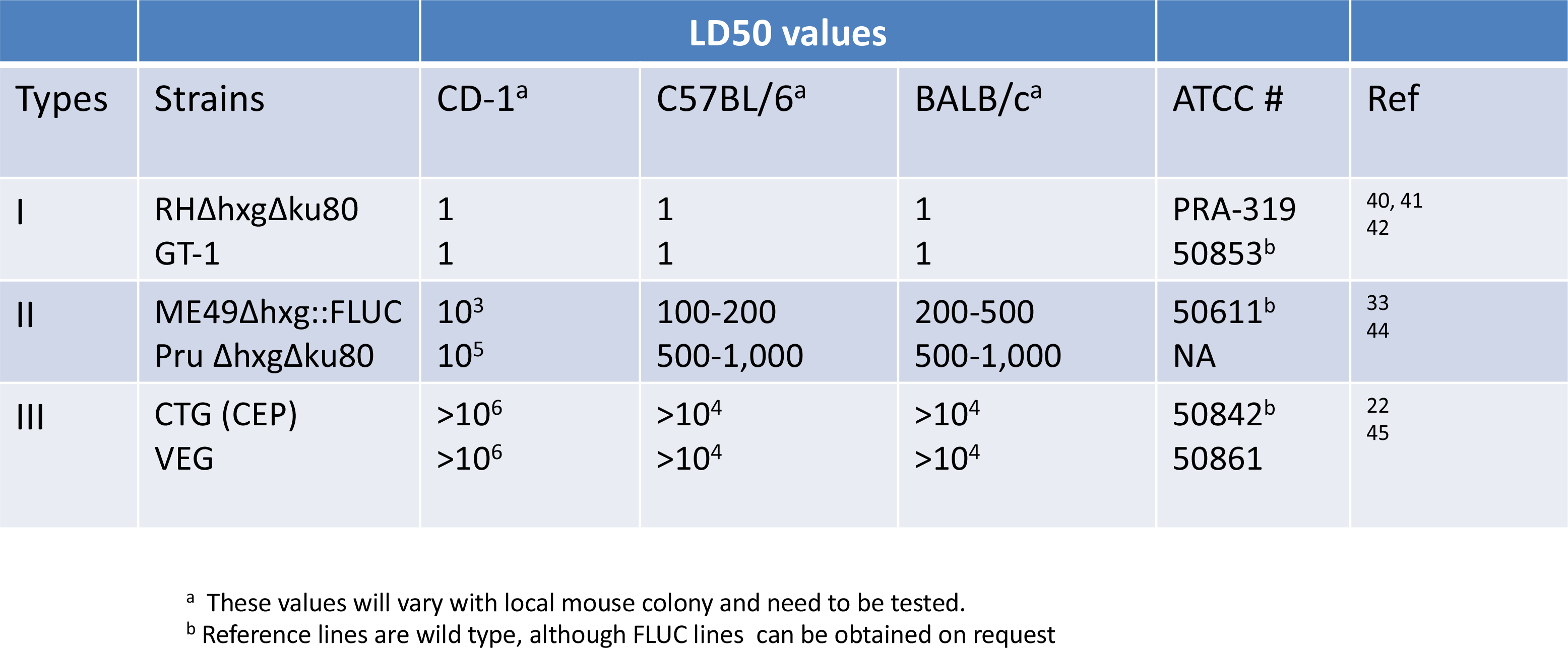
Representative strains of *T. gondii* useful for mouse infection

### 2.2 Challenge studies

1. In vitro tachyzoite cultures of the *T. gondii* strains of interest.
2. Outbred CD-1 and various inbred lines of mice.
3. HHE: 1X Hank’s Balanced Salt Solution, 10 mM HEPES, 1 mM EGTA, sterilized by 0.45 micron filtration.
4. Tuberculin syringes for IP injection.
5. Oral gavage needles 22 g x 1 ½ inch, 2.4 mm tip (Patterson Veterinary, cat. # 07-809-7615).
6. Sterile PBS.
7. Sterile plastic 10 ml syringes equipped with 16, 18, 20 g needles.
8. Sulfadiazine (Thermo Fisher Scientific, cat. # S-6387) solution (0.1 − 0.2 gr/L).

### 2.3 Plaquing assay

1. Tissue culture incubator for maintaining cell cultures at 37 °C, 5% CO_2_ and BSL-2 biosafety cabinet.
2. Human foreskin fibroblasts (HFF) cells (ATCC, cat. # SCRC-1041).
3. D10 medium: For 1 liter, combine 1 package Dulbecco’s Modified Eagle’s Medium (DMEM) powder (Gibco, cat. # 12100046), 3.7 g NaHCO_3_, 100 ml FBS, 10 ml of 200 mM L-glutamine, 1 ml of 10 mg/ml gentamycin, and dH_2_O to 1 liter. Sterilize by 0.45 micron filtration.
4. Tissue culture plates (6 well) containing confluent monolayers of HFF cells.
5. Solution of 1% Crystal violet solution water.
6. Inverted tissue culture microscope, equipped with 4x, 10x, 20x objectives.

### 2.4 ELISA for monitoring infection status

1. High binding ELISA plates (Disposable Sterile ELISA Plates, cat. # 25801).
2. Model 500 Sonic Dismembrator with microprobe (Thermo Fisher Scientific).
3. Tachyzoite culture of *T. gondii*. Most antigens cross-react so RH (Type I) or ME49 (Type II) strains can be used interchangeably for detecting infection with multiple different strain types.
4. Control mouse serum (previously infected positive and non-infected negative animals for reference). Store in aliquots at −80°C until use.
5. Serum samples from infected mice. BD Microtainer tube with serum separator gel (Thermo Fisher Scientific, cat. # 02-675-185). Typically small volumes (10-100 uL) can be collected from saphenous vein or cheek vein puncture. Store in aliquots at −80°C until use.
6. Horse radish peroxidase (HRP)-conjugated secondary antibody (HRP goat anti-mouse IgG) (Thermo Fisher Scientific, cat. # 62-6520).
7. Phosphate buffered saline (PBS).
8. Wash solution: PBS/ 0.05% Tween-20. Plastic squirt bottle for dispensing wash solution.
9. BSA blocking solution: PBS/ 0.05% Tween-20, 1.0% bovine serum albumin (BSA).
10. BSA incubation solution: PBS/ 0.05% Tween-20, 0.1% BSA.
11. Substrate: BD OptEIA Substrate Reagent A, Substrate Reagent B (BD Biosciences, cat. # 51-2606KC).
12. Plate reader for absorbance reading at 450 nm.

### 2.5 Tracking infection by bioluminescence

1. D-Luciferin, potassium salt (Gold Biotechnology, cat # Luck-1G).
2. Isoflurane (Henry Schein Animal Health, cat. # SKU 029405).
3. IVIS Spectrum BL imager (Perkin Elmer) or equivalent instrument capable of detecting bioluminescence.
4. Tuberculin syringes for injection.

### 2.6 Cyst harvesting and staining

1. *Dolichos biflorus* lectin conjugated with FITC (Vector Labs, cat. # FL-1031).
2. Fixating and Permeabilizing Solution: 2X stock consisting of 6% formaldehyde, 0.2% Triton-X-100 in PBS.
3. Blocking solution: 10% normal goat serum in PBS.
4. Glass slides and coverslips.
5. Epifluorescence microscope equipped with phase contrast (10X, 40X) and filter set for detecting FITC.
9. Sterile PBS.
10. Sterile plastic 5, 10 ml syringes equipped with 16, 18, 20 g needles.
6. Polystyrene 15 ml tubes and centrifuge capable of spinning at 400 g.

## 3 Protocols

### 3.1 In vitro propagation of tachyzoites

*T. gondii* are most easily propagated in human foreskin fibroblasts (HFFs) because the host cells reach confluency and stop dividing. These features facilitate passage at high MOI that leads to natural egress at 2-3 day intervals, and also allow for plaque formation on preformed monolayers. Procedures for propagation of HFF monolayers, serial passage of *T. gondii* lines, and harvest of viable tachyzoites have been defined previously (30) and are only briefly summarized here.
1. Passage *T. gondii* tachyzoites by serial passage on HFF monolayers. Typically, strains are inoculated serially using parasites from a freshly egressed culture to inoculate a new monolayer of HFF cells. The flask is inoculated at a high MOI (1:1 or 2:1) to assure uniform infection and host cell lysis in a single round. For Type I stains, natural egress occurs ~2 days after the initial inoculation (or slightly less), while for Type II and III strains it is often 3 days post-inoculation.
2. To passage strains, disperse the contents of a recently egressed culture using a 5 ml pipet to resuspend the culture material and remove cells from the surface (it is also possible to mechanically release as described below).
3. Gently pipet the material using 5 ml pipet to draw the liquid in and out several times to disperse and break up clumps. Dispense parasites using 5-10 drops from a bulb transfer pipet (1 drop ~ 50 μL) to inoculate a new confluent monolayer of HFF cells grown in a T25^2^ flask. For Type I strains, this method results in a 1:10 to 1:20 split by volume (based on a starting volume of 5 mls for culture in a T25^2^ flask) every 2 days. Type II and III strains may require a higher inoculum from 10-20 drops, equating to a 1:10 to 1:5 split by volume. Return the flask to the incubator, 37°C, 5%CO_2_, for 2-3 days.
4. Prior to inoculating mice, it is important to establish the parasites on a consistent passage cycle every 2-3 days, otherwise viability will be compromised.
5. To prepare parasites for inoculation into animals, harvest tachyzoites at the peak of their natural intracellular replication cycle, either at the point of natural egress or shortly before. Any cells remaining on the monolayer at this time point should be heavily infected and will be easily disrupted by gentle pipetting. If necessary, scrape the monolayer to remove cells
6. Resuspend the contents of the flask and passage sequentially through 20, 23, and 25 g blunt needles attached to a 10 ml syringe. When expelling the material from the needle, keep it submerged below the surface to avoid aerating the sample.
7. Filter to separate host cell debris from tachyzoites using 3 micron polycarbonate filter unit. Flush the filter with 10 ml of HHE collecting the flow-through in a 15 ml conical tube.
8. Centrifuge the filtered culture at 400 x *g* for 10 min at 18°C, resuspend the pellet in 10 ml HHE.
9. Count the parasites using a hemocytometer.
10. Dilute the parasites to an appropriate concentration so that injection volumes (typically 0.1 − 0.2 ml) will contain the desired number of parasites.
11. Maintain the parasite at room temperature throughout, chilling them does not result in better viability. Rather it is important to perform these procedures quickly and immediately before you go the facility to inject animals. Different strains of *T. gondii* also vary on how well tachyzoites survive outside of culture, with type I being the most robust. Type II and III strains loose viability much faster and should be used immediately after harvest. We also find the use of Hank’s Balanced Salt Solution based medium is better for resuspending the parasites in compared to PBS, as the former is better for maintaining parasite viability.

### 3.2 Estimating viability by plaquing

1. After returning from the animal facility, use a portion of the unused parasite suspension to perform a plaquing assay and establish the viability of the inoculum.
2. Inoculate 200 − 500 parasites per well of a 6 well plate containing HFF cells growing in DMEM-10% FBS. For a uniform suspension, it is best to place the parasites in 2 ml of D10 medium and then use this to replace the medium that is in each well. Allow parasites to settle by gravity being careful not to swirl the plate (this action creates a vortex that brings the parasites to the center and skews the count). Use triplicate wells per strain.
3. Incubate at 37C, 5% CO_2_ for 7-9 days, depending on the strain. Do not move the plate during this time period.
4. Remove the plate from the incubator, rinse in PBS, and stain with 1% Crystal violet (made in dH_2_O) followed by rinsing in H_2_O.
5. Plaques will appear as clear zone on the stained background. Count by eye or under low power (2-5 X). In the event that plaques have not fully lysed, they can be difficult to visualize. In this case, score foci of infection by examining the stained monolayer under low power using an inverted microscope (5-20 X depending).
6. Expect maximum viability of 50%, however it can also be as low as 5%, especially if tested more than 1-2 hr after initial harvest of the parasites.

### 3.3 Acute virulence model

The following protocol is used to establish “acute virulence” based on serial dilution of parasites and challenge into outbred CD-1 mice. Cumulative survival (or inversely % mortality) is used to evaluate the degree of pathogenicity. This definition of acute virulence has been used to establish the difference between clonal types (11), and map virulence differences between them (12, 24, 31). Additional readouts that are useful to obtain include weight loss, time to death, and tissue burden by bioluminescence (Fig. 1).
1. Harvest tachyzoites from freshly egressed cultures, count, and dilute in HEE as described above.
2. Infect separate groups of 5 CD-1 mice by IP inoculation with serial dilutions of tachyzoite, (i.e. 10, 100 or 1,000 tachyzoites / mouse using Type I parasites). Experiments should be repeated 2-3 times on different days to account for possible variation in the viability of the inoculum.
3. Monitor the animals for weight loss and signs of illness. Use an appropriate end point prior to death, depending on institutional approved protocol.
4. Animals can be monitored for expansion of parasites using luciferase tagged strains and bioluminescence (see **Imaging infection by bioluminescence** protocol).
5. At 30 days post-infection, determine the number of surviving animals. The time point of 30 days is somewhat arbitrary as acutely virulent lineages will generally lead to death prior to day 20.
6. Bleed animals form the saphenous or cheek vein, collecting a small volume (100−200 μL) in a microtainer with serum separator gel. Spin the separator to extract the serum (top) layer from the red cells. Store serum at −20°C before use.
7. Perform ELISA (see **ELISA for monitoring infection** protocol below). to determine the titer in comparison to controls. Seropositivity is an outcome of successful infection. At low inoculum, or low viability of the inoculum, some animals may not become infected and they remain serologically negative. Such non-infected animals are removed from the calculation of % survival.
8. Survival % is calculated as: # infected animals that survive / the number of infected animals (dead animals plus seropositive survivors) × 100. Lower survival equates with higher virulence.

**Figure 1.**
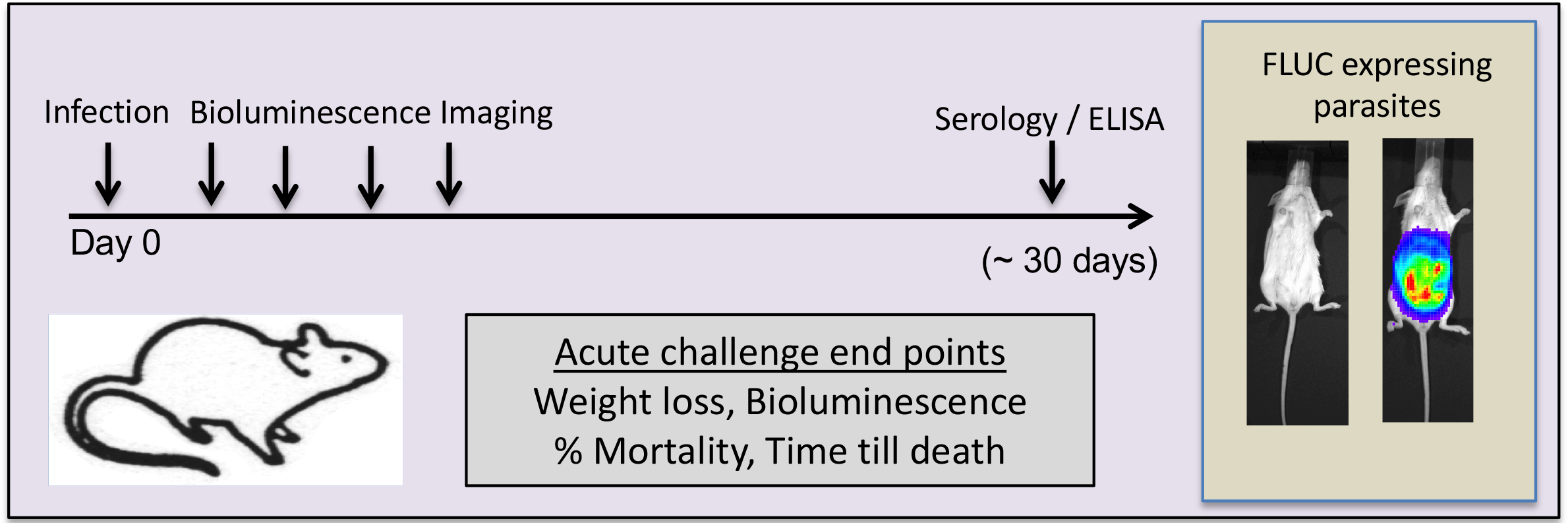
Acute virulence model in outbred CD-1 mice. Following IP challenge with different doses of tachyzoites, mice are followed for 30 days post-infection. Useful end points include weight loss, bioluminescence imaging for luciferase expressing parasites, percent mortality, and time to death. Example shows control and infected mouse imaged for firefly luciferase (FLUC) expression.

### 3.4 Imaging infection by bioluminescence

Bioluminescence provides a powerful way to track infection *in vivo* with the advantage that the same animals can be imaged over time (32). Strains of *T. gondii* have been transfected with various luciferase proteins including firefly (33) and click beetle luciferase (34). These reporters can be imaged by injecting the substrate luciferin *in vivo* just prior to imaging with very sensitive bioluminescence imagers.
1. Resuspend 1 gram of luciferin into 66.7 ml of PBS, filter, dispense in 1.5 ml aliquots, and store in −80°C. One tube should be sufficient for ~5 mice.
2. Weigh mice to determine how much luciferin to inject. Before beginning the experiment, thaw luciferin aliquots.
3. Lightly anesthetize the mice using isoflurane. Inject mice using 10 μL of luciferin (15 mg/ml stock concentration) per gram of weight. For example, if the mouse weighs 25 grams, you will inject 250 μL into the mouse. Injections are performed IP using a tuberculin syringe. Luciferin is readily soluble and rapidly partitions throughout the body, allowing imaging of essentially al tissues, although the penetrance of the signal varies by tissue density.
4. Place the animals in the IVIS chamber using the isoflurane manifold to keep them under light anesthesia to prevent movement during imaging. Image the mice within 20 min of injection for best results.
5. Image the bioluminescence signal following the manufacturers instructions for operation of the IVIS unit.
6. Analyze your data using Living Image ^®^ (Perkin Elmer) and graph results using Prism (Graphpad) or Excel. Appropriate statistical analyses typically involve non-parametric tests using adjustments for multiple comparisons.

### 3.5 ELISA for monitoring infection

#### Antigen preparation

1. Harvest parasites from a freshly egressed culture grown in HFF cells into a 50 ml polystyrene tube. Centrifuge at 400 *g* for 10 min, room temperature. Resuspend the pellet in 10 ml HHE.
2. Count the parasites and resuspend in PBS at a final concentration of 10^8^ cells / ml. Sonicate using 15 sec on/45 sec off cycle for 2 minutes, power 2.8 unit (Fisher Model 500 Sonic Dismembrator with microprobe).
3. Add glycerol to a final concentration of 10%, aliquot, and store aliquots in −80□C.

#### Perform the ELISA

1. Take an aliquot of parasite lysate from the freezer and dilute in PBS (1×10^6^/ ml parasites). Add 100 μl of antigen per well and incubate the plate for 1 hr at 37°C or at 4°C overnight. Cover the plates with lid or parafilm.
2. After incubation, rinse wells 3 times with PBS/ 0.05% Tween-20, using a squirt bottle.
3. Add 250 μl of 1% BSA blocking solution in each well and incubate for 1 hr at room temperature. After incubation, rinse all the wells with PBS/ 0.05% Tween-20 three times.
4. Dilute primary antibody (unknown mouse sera, positive, or negative controls) to appropriate concentration in BSA incubation solution (1:500 − 1:5,000). Add 100 μl of primary antibody to each well and incubate for 1 hr at room temperature. Test each sample and positive and negative controls in duplicate wells. Rinse 3 times with PBS/ 0.05% Tween-20.
5. Dilute HRP-conjugated secondary antibody to 1:2,500 − 1: 10,000 in PBS/ 0.05% Tween-20, 0.1%BSA. Add 150 μl of secondary antibody per well for 1 hr at RT°C. Rinse 4-5 times with PBS/ 0.05% Tween-20. It is important to wash very carefully with PBS after this incubation to avoid unspecific staining because of free peroxidase-conjugated antibodies.
6. Mix equal volume of substrate-A and substrate-B and immediately add 100 μl of this mixture to each well and incubate for 20-30 min at dark. Stop the reaction with 50 μL of 2M H_2_SO_4_ added to each well.
7. Measure the Absorbance at 450 nm with a plate reader.
8. The results of ELISA assays from sample of infected mice are compared to known positive and negative controls. Comparisons can be made by ANOVA to compare the average values of positive and negative controls to individual samples. Alternatively, cutoffs for positive values can be determined as described previously (35).

### 3.6 Chronic tissue cyst bank

Outbred mice are useful to maintain chronic infections that are characterized by low cyst burdens (i.e. 50-200/ animal). Cyst numbers can be amplified in strains of inbred mice that develop higher cysts counts (i.e. 500-3,000 / animal). It is advisable to “bank” strains in CD-1 outbred mice and pass them by sub-inoculating every ~ 6 mos. Cysts are then amplified in susceptible inbred strains (e.g. BALB/c, CBA/J) followed by harvest of tissue cysts over a 1-3 mos period for use in experiments. Although this protocol takes longer to establish, it avoids increases in virulence that can occur with frequent passage and also allows production of high number of cysts for challenge experiments.
1. Inject mice IP with 100-1,000 tachyzoites that have been grown in tissue culture, harvested, purified by filtration, and resuspended in HHE. Viability can be a major issue since it will take you some time to prepare the inoculum and walk to the animal house. Use only freshly egressed parasites that have been grown under optimum conditions (i.e. 2-3 day synchronous cultures).
2. Intermediately virulent strains (i.e. ME49) have LD_50_s around 10^3^−10^5^ in outbred mice. Therefore it is usually not necessary to treat the mice to prevent death. You want the mice to be heavily infected without dying. Use animals that survive the highest dose possible to obtain higher cyst counts.
3. Virulent Type I strains like GT-1 are more problematic as they will invariably cause death even at low doses. Use an inoculum of 50-100 tachyzoites IP (at lower doses some mice will not become infected).
4. To prevent accidental death with high virulence strains, it is necessary to treat the mice with sulfadiazine (dissolved in the drinking water). Begin treatment with a dose of 0.4-0.5 g/L on day 3 or 4 and treat for 6-10 days or until they recover. Even for less virulent strains (i.e. ME49) it is sometimes necessary to treat with sulfadiazine at lower doses (0.1 or 0.2 g/L as above).
5. Harvest the brain of chronically infected mice at 1-3 months post-infection and determine the cyst burden by homogenizing and counting a fraction of the homogenate (see **Cyst harvesting and staining** protocol below). It is possible to obtain a sufficient number of cysts from a single animal to use in subsequent challenge studies, depending in the number of animals and desired inoculum.
6. To maintain chronic cysts, establish a bank of CD-1 mice and passage them at 6 mos intervals. Infected animals should be humanely sacrificed, brains removed and homogenized. After staining and counting a proportion of the brain (see **Cyst harvesting and staining** protocol below) serial passages are done by oral gavage of 5-10 cysts per animal into naïve mice.
7. To expand cyst numbers, inoculate 5-10 cysts IP or PO into inbred CBA/J or BALB/c mice, which are more susceptible and will lead to higher cysts counts. Infections can be passed sequentially for 2-3 times in inbred mice, but do not use this for long-term passage as the strain will increase in virulence.

### 3.7 Cyst harvesting and staining

This protocol is designed to help visualize tissue cysts that form in the brain of chronically infected animals. Tissue cysts can be recognized by their appearance in phase contrast microscopy (refractive, slightly amber cyst wall with internal granular material). However, it is much easier to identify them based on positive staining with FITC-conjugated *Dolichos biflorus* lectin (DBL) (36).
1. Sacrifice chronically infected animals using an approved method.
2. Remove the brain and place in a sterile conical 15 ml tube containing ~ 2 ml PBS. Gently mince the brain using a 16 g needle. Draw the tissue into a 5 ml syringe and gently expel. Repeat 3 times using 16 and 18 g needles to homogenize the tissue.
3. Add an aliquot of brain lysate (typically ¼ of the total or 0.5 ml) to an equal volume of Fixing and Permeabilizing Solution in a 15 ml polystyrene centrifuge tube. Incubate for 20 min at 4°C.
4. Spin at 400 g, 4°C, 5 min. Resuspend in 4 ml PBS/10% goat serum. Spin down pellet at 400 g for 5 min and remove supernatant.
5. Retain the pellet and add 1 ml of 10% goat serum in PBS, containing 2~4 μL of FITC-conjugated-lectin stock solution. Incubate at room temperature for 45-60 min.
6. Wash twice by centrifugation at 400 g, 15°C for 5 min. Resuspend the pellet each time in 4 ml PBS + 10% goat serum.
7. After the final wash, re-suspend cysts in PBS at the original volume of tissue homogenate used.
8. Add 12.5 μL of sample to a microscope slide, cover the sample with coverslip, and screen under fluorescence microscope using a 10X objective. Scan the slide at 10X and when you see a possible cyst, confirm by examining at 40X. Measure the diameter of the cysts using a calibrated ocular micrometer. Scan the entire slide and determine the cyst number per 12.5 μL. For each sample, count 4 aliquots and determine the total number of cysts. Calculate the number of cysts per brain (total number of cysts in 4 aliquots (50 μL) x 40 = total number per brain).

### 3.6 Chronic infection model

There are many advantages to working with less virulent strains that readily produce chronic infections in mice as both the acute and chronic phases can be studied (Figure 2). Type II strains, such as ME49 (popular clones include the B7 clone, and PTG) and Pru, provide convenient lab-adapted strains for this purpose. Susceptibility of different mouse strains to infection with Type II strain parasites varies in large part due to differences in MHC with BALB/c (H-2d) and C3H/HeN (H-2k) mice being more resistant than C57Bl/6 (H-2b) (37). Hence, the combination of Type II strains with different mouse backgrounds can be used to study acute virulence, chronicity and reactivation. Genetic mutants in the Type II strain are particularly useful for studying attenuation as they are more likely to reveal partial phenotypes that may be masked by the extremely high virulence of Type I strains.
1. Inject mice IP with tachyzoites that have been grown in tissue culture, harvested, and resuspended in HHE. Refer to Table 1 for approximate doses depending on the mouse strain being used. Viability can be a major issue since it will take you some time to prepare the inoculum and walk to the animal house. Use only freshly egressed parasites that have been grown under optimum conditions (i.e. 2-3 day synchronous cultures).
2. When testing genetic mutants vs. wild type, it is useful to bracket the inoculum starting at the LD_50_ and increased by half log or log intervals. Typically 5 mice are used per group. It is generally necessary to repeat the experiment at least once since the outcome is influenced by differences in viability. Statistical analysis of Kaplan Meier survival curves is readily performed in Prism (GraphPad).
3. Plaque an aliquot of the parasite suspension after injecting them to assure that the different parasite isolates had approximately equal viability.
4. During the acute phase (day 4-20) monitor expansion of the parasites using the **Imaging infection by bioluminescence** protocol. It is also useful to monitor weight loss and regain as the mice recover following acute infection.
5. Animals are generally followed for 30-60 days, and percent survival is calculated as above. In most cases lethal outcomes will be apparent by day 20, although in rare occasions mutants will show delayed kinetics of death. Use an appropriate end point prior to death, depending on institutionally approved protocol.
6. At the end of 30-60 days, sacrifice the animals using an approved method. Remove the brain and perform the **Cyst harvesting and staining protocol.** Statistical analyses of tissue cyst numbers is usually performed using non-parametric tests with adjustment for multiple comparisons.
7. It is also possible to test the oral infectivity of tissue cysts by oral gavage (or IP injection) into naive animals. Administer tissue cysts in 100-200 μL PBS suspension, inoculating a dose of 5-10 cysts per animal. Progression of infection can be monitored by sero-conversion using the **ELISA** protocol.

**Figure 2.**
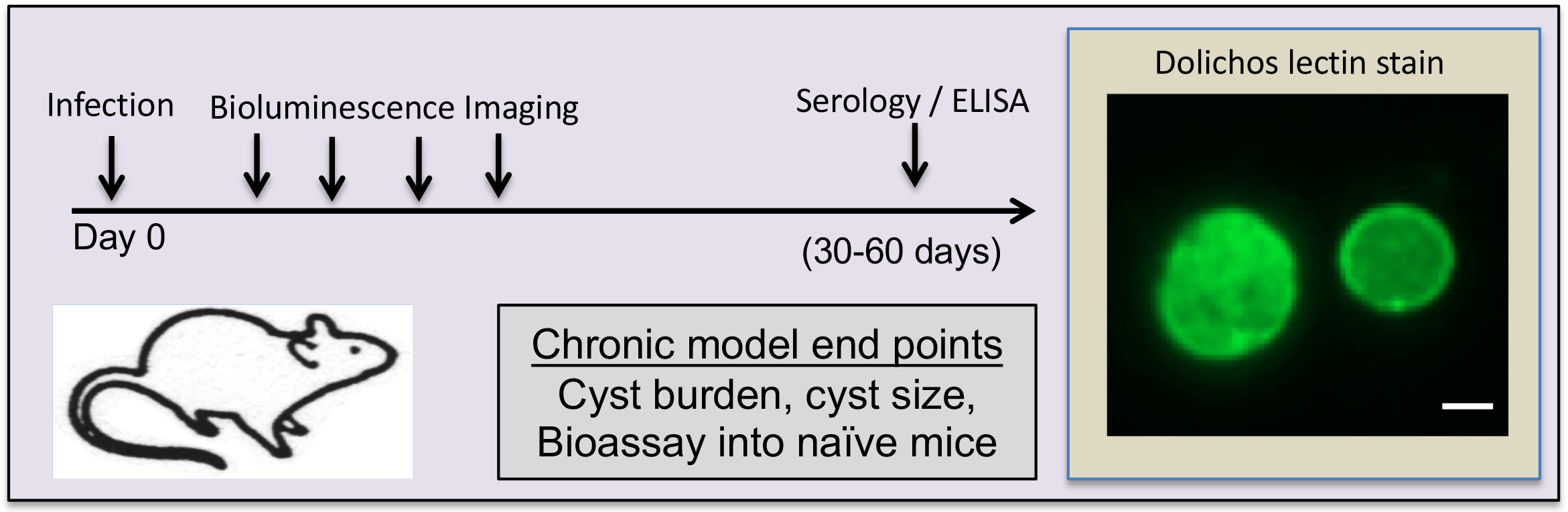
Chronic infection model in inbred mice. Infections can be administered either by IP injection of tachyzoites or by oral feeding of tissue cysts derived from chronically infected animals. Useful end points include dose-dependent mortality, cyst burden, and cyst size. It is also possible to use bioluminescence imaging to evaluate differences in parasite numbers during the acute phase. The infectivity of tissue cysts can also be tested by sub-inoculation into naive animals. Example shows tissue cyst stained with *Dolichos biflorus* lectin. Scale bar = 10 microns.

## 4 Notes

### Strain origins and derivatives

The type I RH strain was originally isolated from an adolescent who succumbed to fatal encephalitis (38). This lab-adapted strain has been modified to express various transgene reporters including luciferase for bioluminescence imaging (32). One isolate commonly used for genetic studies has been modified to delete the hypoxanthine, xanthine, guanosine phosphoribosyl transferase gene (*HXGPRT* also known as *HXG*), allowing for positive and negative selection (39), as well as disruption of the *KU80* gene, which results in increased levels of homologous recombination (40, 41). The RHΔ*hxg*Δ*ku80* strain maintains the full virulence of the wild type RH strain. Another commonly used Type I strain is GT-1, which was isolated from a goat (42), and which maintains the entire life cycle unlike the commonly used RH strain (12). Type II strains are also widely used for creating chronic infections and for virulence studies where the LD_50_ can be titrated in inbred mice. The most common of these is the ME49 strain, originally isolated from a sheep in California (43): it readily generates chronic infection in mice and undergoes the entire life cycle in cats (23). The ME49 strain is also available as a Δ*hxgprt* knockout for facilitating selection and is tagged with firefly luciferase (FLUC) (33). Another commonly used Type II strain is Prugniaud (aka Pru), originally isolated from a congenital human infection in France, and including a variant that is lacking both *KU80* and *HXGPRT* (PruΔ*hxg*Δ*ku80)* (44). Although less commonly used, Type III strains also generate chronic infections in mice but are rarely used for infection studies due to their relative avirulence (4). The Type III strain CTG was originally isolated from a cat in New Hampshire (22) and the VEG strain was isolated from an immunocompromised patient (45).

### Acute virulence

Infection with Type I strains is always lethal in laboratory mice, and any animals surviving at low inoculum are sero-negative (i.e., never infected). We favor the use of outbred mice, which are more resistant, as they clearly discriminate from the high virulence of Type I strains in comparison to strains that have intermediate virulence that depends on the mouse strain. This definition of acute virulence has been used to establish the difference between clonal types (11), and map virulence differences between them (12, 24, 31). With Type II strains, the LD_50_ in outbred and inbred mice differs substantially (Table 1) and aspects of pathogenesis during chronic infection can be evaluated in surviving mice. The LD_50_ of Type II strains is also somewhat dependent on the local mouse colony, so it needs to be titrated in each instance.

### Bioluminescence

The sensitivity of bioluminescence detection *in vivo* is not easy to relate directly to parasite number and it is likely that high tissue burdens (i.e. > 10^5^ parasites / gram) are needed to detect infection. Additionally, it can be more challenging to detect infection in deep tissues in particular in the CNS due to the fact that the cranium partially blocks the signal. Nonetheless, this method has proven highly useful for tracking acute infection and dissemination (8, 31, 46), as well as reactivation of infection in immunocompromised mice (47).

### Alternatives for monitoring tissue burdens

Other approaches for monitoring the number of parasites in tissue following acute or chronic infection have been developed based on PCR (48) or plaquing (13). These methods have the advantage of being quantitative in terms of relating to genome equivalents or infectious units. However, PCR can over-estimate the number of parasites due to remnant DNA that is not derived from viable organisms (48). Using an RNA target and performing qRT-PCR can partially compensate for this possibility (49), although neither method measures viable parasites. The main limitations of these methods are they are time consuming, and do not allow the possibility of sampling the same mouse repeatedly.

## Acknowledgements

We thank many former members of the laboratory for developing and refining these protocols over the years, including Mike Behnke, Kevin Brown, Ildiko Dunay, Blima Fux, Dan Howe, Asis Khan, Dana Mordue, and Chunlei Su. We are also grateful to Christopher Hunter and Yasu Suzuki for helpful advice on animal infection models, and to David Bzik, Vern Carruthers, J.P. Dubey, and Laura Knoll for generously providing strains. Supported in part by NIH grants AI118426 and AI034036 to LDS.

